# MYC Dysregulates Mitotic Spindle Function Creating a Dependency on TPX2

**DOI:** 10.1101/272336

**Authors:** Julia Rohrberg, Alexandra Corella, Moufida Taileb, Seda Kilinc, Marie-Lena Jokisch, Roman Camarda, Alicia Zhou, Sanjeev Balakrishnan, Aaron N. Chang, Andrei Goga

## Abstract

The MYC oncogene promotes tumorigenesis in part by facilitating cell cycle entry thus driving cellular proliferation. Tumors that overexpress MYC frequently demonstrate aneuploidy, numerical chromosome alterations associated with highly aggressive cancers, rapid tumor evolution, and poor patient outcome. While the role of MYC in overcoming the G1/S checkpoint is well established, it remains poorly understood whether MYC induces chromosomal instability (CIN). Here, we identify a direct influence of MYC on mitotic progression. MYC overexpression induces defects in microtubule nucleation and spindle assembly promoting chromosome segregation defects, micronuclei and CIN. We examined which mitotic regulators are required for the survival of MYC-overexpressing cells and found a reliance on high TPX2 expression. TPX2, a master microtubule regulator, is overexpressed together with MYC in multiple cell lines, in mouse tumor models and in aggressive human breast cancers. High TPX2 expression is permissive for mitotic spindle assembly and chromosome segregation in cells with deregulated MYC, whereas TPX2 depletion blocks mitotic progression, induces cell death and prevents tumor growth. Importantly, attenuation of MYC expression reverses the mitotic defects observed, even in established tumor cell lines, implicating an ongoing role for high MYC in the persistence of a CIN phenotype in tumors. Here, we implicate the MYC oncogene as a regulator of spindle assembly and dynamics and identify a new MYC-TPX2 synthetic-lethal interaction that could represent a future therapeutic strategy in MYC-overexpressing cancers. Our studies suggest that blocking MYC activity can attenuate the emergence of CIN and tumor evolution.

## Introduction

Aneuploidy has been implicated in tumorigenesis for decades with more than 70% of common solid tumors found to be aneuploid^1,2^. Aneuploidy, which is a state of abnormal chromosome number, is frequently caused by chromosomal instability (CIN), chromosome missegregation that leads to chromosome loss or gain following mitotic division^3,4^. CIN is a major driver of tumor evolution and promotes drug resistance and metastasis^5–7^, however the major mechanisms that mediate CIN in most common tumors remains poorly understood, since mutations in spindle-associated genes occurs infrequently^8–12^.

The MYC oncogene is frequently overexpressed in a wide variety of aggressive and metastatic tumors and has been associated with structural chromosomal aberrations and aneuploidy^13–16^. However, whether MYC contributes directly to aneuploidy, possibly through the induction of CIN, is unknown. One of the key biological functions of MYC is its ability to facilitate entry and progression through G1 and S phases of the cell cycle by regulating gene transcription^17^. Whether MYC also impacts mitotic progression is unclear. Cells with elevated MYC activity are sensitive to mitotic interruption such as treatment with microtubule targeting agents, mitotic kinase inhibitors or siRNA against spindle-related genes^18–25^. However, a molecular mechanism for MYC’s synthetic-lethal interactions with mitotic regulators is missing. Clarifying such a mechanism could reveal novel treatment strategies for aggressive cancers.

CIN is prevented by the spindle assembly checkpoint (SAC) that detects erroneous microtubule-kinetochore attachments ensuring faithful chromosome segregation^26^. Since in human cancers impairments of the SAC are rarely detected, merotelic attachments are believed to be a major route to CIN^4,27,28^. A single merotelic kinetochore is attached to both spindle poles thus satisfying the SAC^1^. Various defects in spindle morphology and assembly can cause an increase in merotelic attachments in cancer cells leading to CIN^1,29,30^. One key player of spindle formation is the microtubule binding protein TPX2. TPX2 is required for the initiation of microtubule growth from chromosomes and controls spindle scaling, centrosome movement and chromosome segregation^31–33^. TPX2 overexpression is highly correlated with CIN and is found in a many aggressive human tumors, but the precise role of TPX2 for CIN formation and maintenance remains unclear^34,35^.

Here, we investigate whether MYC directly induces CIN in various cellular models. We use gene-expression data and siRNA screening approaches to identify mitotic regulators that are important for the survival of cells with deregulated MYC.

## Methods

### Lentiviral constructs, virus production and infection

The FUCCI plasmid pCDII-EF-MCS containing mKO2-hCdt1(30/120) and mAG-hGeminin(1/110) have been described previously^55^ (GenBank accession number AB370332 and AB370333). H2B-mCherry pLenti6/V5-DEST has been described previously^77^. For inducible shRNA expression the following short hairpin sequences were cloned into the Tet-PLKO-puro vector^78^: *shTPX2* no. 1: 5’-CCGGGAACAATCCATTCCGTCAAATCTCGAGATTTGACGGAATGGATTGTTCTTTTT −3’; *shTPX2* no. 2: 5’-CCGGCTAATCTTCAGCAAGCTATTGCTCGAGCAATAGCTTGCTGAAGATTAGTTTTT −3’. For control shRNA against GFP with the following sequence was used: 5’-CCGGTACAACAGCCACAACGTCTATCTCGACATAGACGTTGTGGCTGTTGTATTTTTG −3′. 293T cells used for producing lentiviral particles and were grown in DMEM medium with high glucose supplemented with 10% FBS and penicillin/streptomycin. Cells were infected with lentiviral constructs and selected for 5 days with 0.5 μg/ml puromycin starting 48 h after infection. FUCCI expressing cells were sorted using fluorescence associated cell sorting on a BD FACSAria II flow cytometer (Becton Dickinson).

### Cell lines and propagation

All breast cancer cell lines were purchased from ATCC and cultured as previously described^79^. RPE-NEO and RPE-MYC were a gift from Michael J Bishop and cultured as previously described^22^. RPE-NEO were not used beyond passage 16. Primary human mammary epithelial cells expressing MYC-ER and shRNA specific for the p16 isoform-encoding sequence of *CDKN2A* (HMEC) were generated and cultured as previously described^37^. Although expression of p16 shRNA delays senescence, the cells are not immortalized and undergo spontaneous senescence when continuously cultured. Therefore, these cells were not used beyond 12 passages after their derivation. HMEC cells were treated with 500 nM 4-hydroxytamoxifen (TAM) to induce MYC activation for 48 hours. All cell lines were continuously tested negative for mycoplasma contamination using PCR.

### RNAi knock down

All siRNAs were purchased from GE Dharmacon (ON-TARGETplus, four siRNAs per gene). For Figure 4a 50 nM siRNA was used. For any other TPX2 knock down experiment 1.7 nM siRNA was used. For MYC knock down 30 nM siRNA was used (Figure 1h). Cells were transfected using Lipofectamine RNAiMAX Transfection Reagent (Thermo Fisher), according to the manufacturer’s instructions.

**Figure 1.**
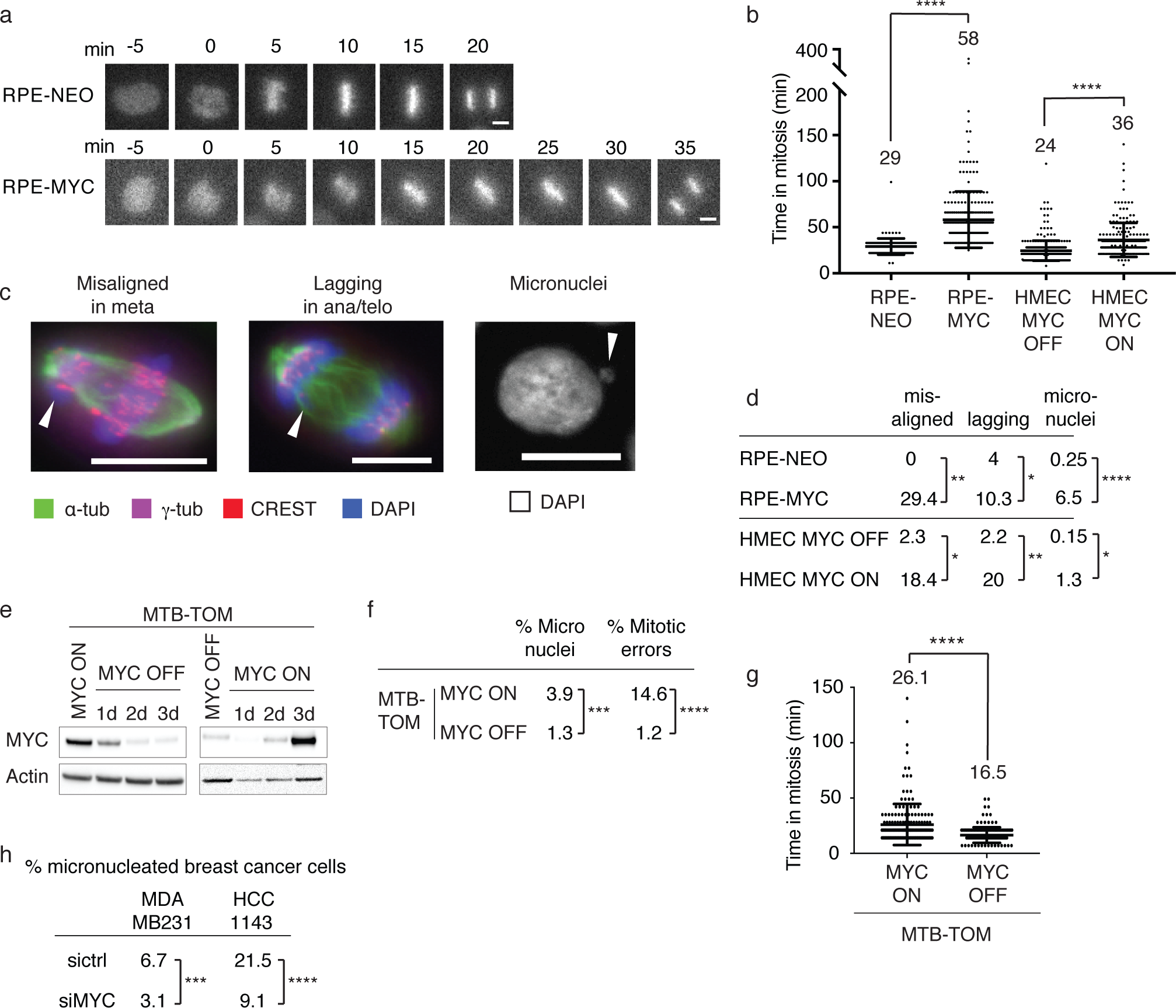
MYC reversibly induces CIN. **a** Representative images of fluorescent time-lapse microscopy of RPE-NEO and RPE-MYC cells expressing H2BmCherry. Pictures were taken every 5 min. Scale bar, 10 μm. **b** Quantification of time from chromosome condensation to anaphase onset in RPE-NEO, RPE-MYC, HMECs expressing MYC-ER in the absence (MYC OFF) and presence (MYC ON) of MYC activation for 3 days. Mean +-S.D., t-test, n=140-380. **** p<0.0001. **c** Representative images of misaligned chromosomes in metaphase, lagging chromosomes in ana- or telophase and micronuclei in RPE-MYC cells. Fixed cells were stained with anti-δ-tubulin (spindle), anti-8-tubulin (centrosome), anti-CREST (centromere) and DAPI (DNA). Scale bar, 10 μm. **d** Quantification of misaligned chromosomes in metaphase, lagging chromosomes in ana- or telophase and micronuclei in RPE-NEO, RPE-MYC and HMECs expressing MYC-ER in the absence (MYC OFF) and presence (MYC ON) of MYC activation for 3 days. Fisher’s exact test, n=100-300 mitotic cells and n=800-1000 cells for micronuclei. * p<0.05, ** p< 0.01, **** p<0.0001. **e** Western blot of MYC in a cell line derived from an MTB-TOM tumor expressing a doxycycline (dox)-inducible MYC construct. Cells were grown in presence of dox (MYC ON) and dox washed out (MYC OFF) and added back for the indicated time. **f** Percent MTB-TOM cells with micronuclei and mitotic errors in the presence (MYC ON) and absence (MYC OFF) of MYC expression for 4 days. For micronuclei quantification, cells were fixed and stained with DAPI. For quantification of mitotic errors, cells with lagging chromosomes and chromatin bridges were counted from time-lapse microscopy experiments. Fisher’s exact test, n=1628 and 520 for micronuclei and 178 and 164 for mitotic errors. *** p<0.001, **** p<0.0001. **g** Quantification of time from chromosome condensation to anaphase onset from time-lapse fluorescent microscopy of MTB-TOM cells expressing H2BmCherry in the presence (MYC ON) or absence (MYC OFF) of MYC expression for 4 days. Mean +-S.D. t-test, n=178 and 164. ****p<0.0001. **h** Percent of micronucleated triple-negative breast cancer cell lines MDAMB231 and HCC1143 3 days after transfection with control (sictrl) and MYC (siMYC) siRNA. Cells were fixed and stained with DAPI. Fisher’s exact test, n=628-1056. *** p<0.001, **** p<0.0001.

### Immunofluorescence microscopy

Cells were grown on glass coverslips (Fisherbrand), fixed with 100% methanol at −20 °C for 3 min and stained with 4’,6-diamidino-2-phenylindole (DAPI, 1:10,000) to visualize DNA and antibodies against: anti-㎱-tubulin (1:1,000, DM1 ㎱, T6199, Sigma), anti-centromere (CREST, 1:50, 15-234 Antibodies Incorporated), anti-δ-tubulin (1:1,000, T3559, Sigma), anti-TPX2 (1:1,000, HPA005487, Sigma). Secondary antibodies conjugated to Alexa Fluor-488 (1:1,000, A11029, Thermo Fisher), Alexa Fluor-594 (1:1,000, A11014, Thermo Fisher) and Alexa Fluor-647 (goat, 1:1,000, ab150079) were used. Fixed cells were imaged with exposure times of 5-200ms with DAPI, GFP, TRITC, and CY5 filter cubes and a mercury arc lamp on a Zeiss AxioPlan2 epifluorescence microscope (operated by MicroManager 1.4.13) with a 40x 1.3 DIC oil objective and a QIClick camera (QImaging). Images were recorded with a *Z*-optical spacing of 0.2 μm and analyzed using the Fiji software^80^. Centrosome distance was measured in cells with correctly aligned metaphase plates and in which the centrosomes were found within the same plane. TPX2 immunofluorescence (Supplementary figure 4) was imaged at 37°C with a 100x NA 1.49 objective lens (CFI APO TIRF; Nikon) on an inverted microscope system (TE2000 Perfect Focus System; Nikon) equipped with a Borealis modified spinning disk confocal unit (CSU10; Yokogawa) with 200-mW, 405 nm, 488 nm, 561 nm and 643 nm solid-state lasers (LMM5; Spectral Applied Research), electronic shutters, a Clara cooled scientific-grade interline CCD camera (Andor), and controlled by NIS-Elements software (Nikon).

### Live-cell imaging

For live-cell analyses, cells expressing H2B-mCherry or FUCCI were seeded into 12 well plates at 50,000 cells/well and followed by time-lapse microscopy at 37 °C and 5% CO_2_. Images were acquired on an inverted microscope (Nikon Eclipse Ti), operated by NIS-Elements software and equipped with a CoolSNAP HQ2 CCD camera (Photometrics). Cells were imaged with phase contrast (20-100 ms exposure), 488 nm and 594 nm laser light (75-200 ms exposure) through a 20x 0.45 Ph1 objective using perfect focus every 7-10 min, in a stage-top incubation chamber (Okolab) maintained at 37°C and 5% CO_2_. The time from nuclear envelope breakdown (loss of nuclear FUCCI) or chromosome condensation (H2B-mCherry) until the beginning of anaphase (start of chromosome movement to the poles) or cell death was determined. Graphs were generated using the Prism software package, version 7 (GraphPad Software).

### Microtubule regrowth assay

Cells were seeded onto coverslips into 24-well plates at 100,000 cells per well. To depolymerize microtubules cells were incubated with 0.2 μg/ml nocodazole for 5 hours. Coverslips were drained and placed into media without nocodazole. Cells were fixed and stained after the indicated time points. Microtubule asters at chromatin and centrosomes were counted and the distance between centrosomes measured. For quantifying cells with aligned chromosomes, all cells that started to align their chromosomes plus cells with metaphase plates were included.

### Western blotting

Cells were lysed in Laemmli buffer (60 mM Tris-HCl, pH 6.8, 1 μM DTT, 2% (w/v) SDS) supplemented with protease inhibitor cocktail (Roche) and phosphatase inhibitor cocktail (Roche). Protein concentration was determined using the DC Protein Assay (BioRad). 30 μg Protein extracts were resolved using 4-12% SDS-PAGE gels (Life Technologies) and transferred to nitrocellulose membranes using iBlot (Life Technologies). Membranes were probed with primary antibodies overnight on a 4 °C shaker, then incubated with horseradish peroxidase (HRP)-conjugated secondary antibodies, and signals were visualized with ECL (Bio-Rad). The following primary antibodies were used: Anti-β-actin (actin) (1:10,000, sc-47778 HRP, Santa Cruz), Anti-c-MYC (MYC) (1:1,000, ab32072, Abcam), Anti-TPX2 (1:1,000, HPA005487, Sigma), Anti-cleaved PARP (1:1000, 9542, Cell signaling)

### Cell viability assays

To screen spindle genes in RPE-NEO and RPE-MYC cells were seeded in 12-well plates at 50,000 cells/well and transfected with 50 nM siRNA as described above or treated with 10 μM of the CDK1 inhibitor Purvalanol (Figure 4a). Cells were harvested after 72 h and cell viability was assessed by performing the flow cytometry-based Guava ViaCount viability assay (Millipore) according to the manufacturer’s instructions. For further validation of the effects of siTPX2 treatment, cells were seeded in 6-well plates at 100,000 cells per well and transfected with 1.7 nM siRNA as described above. Cells were harvested at 72 h and cell viability determined using the PrestoBlue cell viability reagent (Thermo Fisher) according to the manufacturer’s instructions.

**Figure 4.**
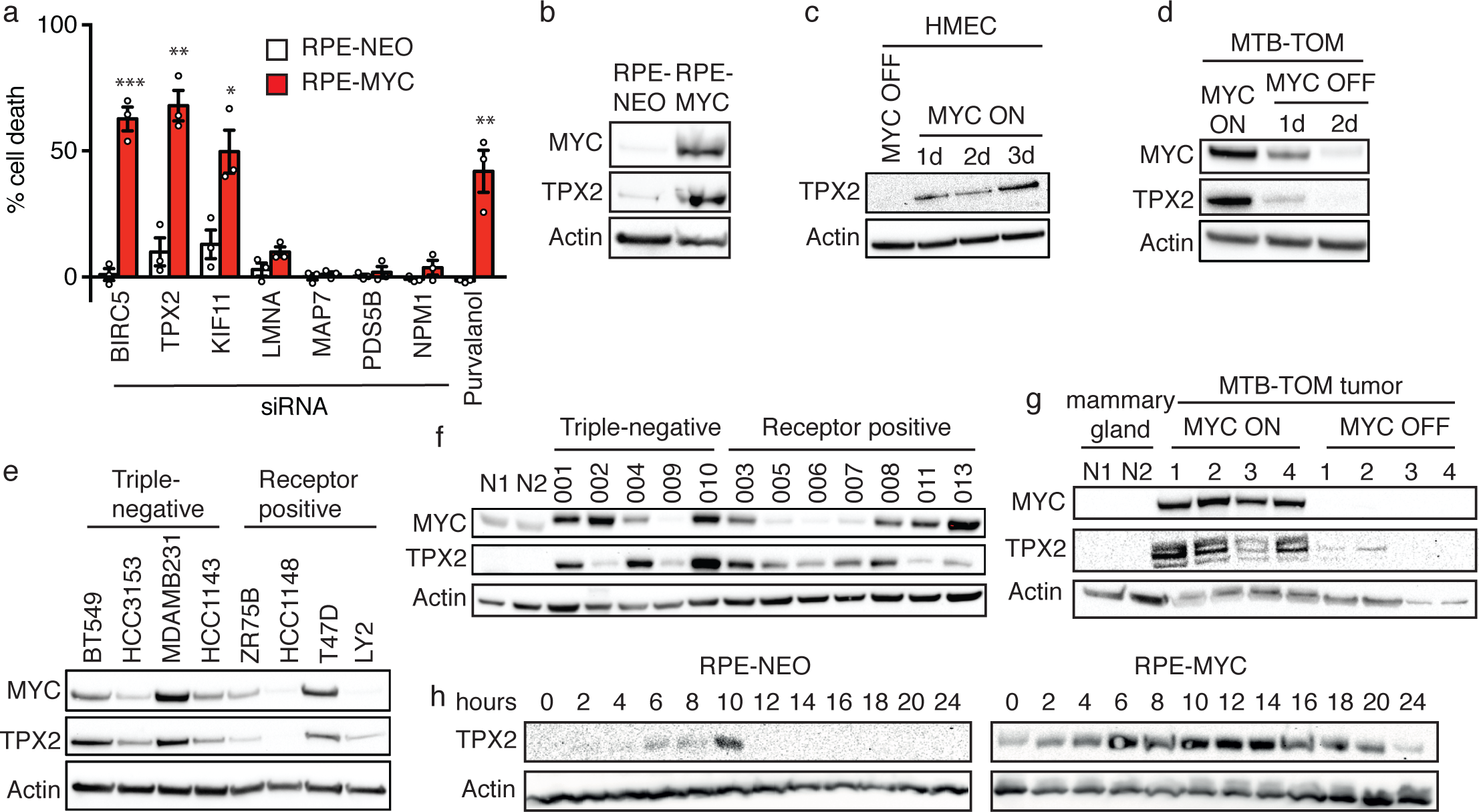
TPX2 expression correlates with MYC. **a** Percent cell death in RPE-NEO (white bars) and RPE-MYC (red bars) cells 3 days after siRNA knock down of spindle genes normalized to control siRNA, or treatment with 10 ¼M Purvalanol A normalized to DMSO. Bars, mean +-S.E.M., t-test, n=3. * p<0.05, ** p< 0.01, *** p<0.001. **b** Western blot of MYC and TPX2 in RPE-NEO and RPE-MYC. **c** Western blot of TPX2 in HMECs expressing MYC-ER in presence (MYC ON) or absence of MYC activation (MYC OFF) for the indicated time points. **d** Western blot of MYC and TPX2 in MTB-TOM cells in presence (MYC ON) and absence (MYC OFF) of doxycycline. **e** Western blot of MYC and TPX2 in triple-negative, high MYC, and receptor positive, low MYC, breast cancer cell lines. **f** Western blot of MYC and TPX2 in triple-negative and receptor positive human patient-derived xenograft tumors and normal human mammary gland (N1 and N2). **g** Westernblot of MYC in mouse mammary gland (n=2), MTB-TOM tumors (MYC ON, n=4) and MTB-TOM tumors from mice that were taken off doxycycline diet for 3 days to switch off MYC expression (MYC OFF, n=4). **h** Westernblot of TPX2 in RPE-NEO and RPE-MYC after release from a double thymidine block to arrest cells in S phase at the indicated time points. Mitotic entry was observed in both cell types ~ 10 hrs after release.

### Cell cycle profiles

For determining cell cycle profiles after siTPX2 treatment cells were seeded in 6-well plates at 100,000 cells per well and transfected with 1.7 nM siRNA as described above. Cells were harvested at 72 h, fixed in 95% ethanol overnight at 4 °C and incubated in 80 μg/ml propidium iodide 150 μg/ml and RNaseA in PBS for 1 hour. Flow cytometry was performed on a BD LSRII flow cytometer (Becton Dickinson) and 20,000 events were counted. Data were analyzed using FlowJo 2 (FlowJo, LLC).

### Animal studies

All protocols described in this section regarding mouse studies were approved by the UCSF Institutional Animal Care and Use Committee. The study complied with all relevant ethical regulations. The ethical end point for tumor experiments was reached when a tumor reached ≥2 cm in any single dimension.

### MTB-TOM tumor generation and immortalization of cell line

MTB-TOM (MMTV-rtTA/TetO-*MYC*) mice were generated as previously described^39^. Mice were bred and maintained off of doxycycline. At 12-15 weeks of age, female mice were put on doxycycline (200 mg/kg doxy chow, Bio-Serv) to induce MYC expression and tumorigenesis. Mice were monitored daily for tumor growth by inspection and caliper measurement in two dimensions. Mice were put off doxycycline when tumors reached 1 cm in any single dimension for 3 days to switch off MYC expression. Tumors and mammary glands were flash-frozen in liquid nitrogen.

To generate a cell line, a tumor from an MTB-TOM mouse was removed, weight determined, chopped and incubated in 5 ml/g of tissue of RPMI 1640 supplemented with 10 mM HEPES pH 7.4, 2.5% FBS, 1 μg/ml doxycycline and 1 mg/ml collagenase IV (Sigma) for 1 h at 37°C and 200 rpm shaking. The tissue was washed 3 times with RPMI 1640 supplemented with 10 mM HEPES and 2.5% FBS and plated into PyMT medium (DMEM/F12 supplemented with 5 pg/ml Insulin, 1 pg/ml Hydrocortisone, 10 ng/ml EGF, 10% FBS, 10 U/ml penicillin, 10 mg/ml streptomycin, 50 pg/ml Gentamycin, 2 mM Glutamine, 10 mM HEPES pH 7.4, 1 ¼g/ml doxycycline) with centrifugation at 1200 rpm for 2 min and 3 quick spins (1200 rpm for 5 s). 1×10^6^ cells were passaged every 3 days. After passage 10 cells were grown in DMEM supplemented with 1 μg/ml doxycycline for another 10 passages. To switch off MYC expression cells were grown in the absence of doxycycline for at least 3 days and not more than 7 days.

### Patient-derived xenograft models

All human samples used to generate PDX tumors, as well as the human non-tumor samples, were previously described^81^.

### Mice xenograft studies

To test the influence of shTPX2 expression HCC1143 and BT549 cells were infected with Tet-pLKO-puro-shTPX2 no 1. 1 x 10^6^ cells were transplanted into the cleared mammary fat pads of 4-week-old female NOD/SCID mice (Taconic Biosciences). After the tumors reached 1 cm^3^, mice were fed doxycycline chow (200 mg/kg, Bio-Serv) to induce *shTPX2* or *shGFP* expression. Tumor growth was monitored 3 times a week by caliper measurement in two dimensions. Mice were euthanized after 30 d of treatment or after tumors reached 2 cm in any dimension.

### RNAseq and gene expression analysis

For RNAseq and gene expression analysis, 11 MTBTOM tumors, 3 mammary glands from mice that were never fed doxycycline and 5 tumors from mice that were taken off doxycycline for 72 hours were flash-frozen in liquid nitrogen. 3 biological replicates of HMECs expressing a MYC-ER fusion construct were grown in presence of 0.5 uM tamoxifen to activate MYC or solvent for 72 hours. RNA was isolated using the RNAeasy kit (Qiagen). Library preparation and Illumina RNAseq was performed by Q^2^ Solutions (www.q2labsolutions.com).

Differential gene expression analysis of HMEC RNAseq data were processed using the ArrayStudio software (Omicsoft). Differential gene expression analysis of MTBTOM tumors was processed using the *limma* R package (Ritchie et al., 2015). TCGA breast-invasive carcinoma data set was sourced from data generated by TCGA Research Network (http://cancergenome.nih.gov), made available on the University of California, Santa Cruz (UCSC) Cancer Browser. Genes that were significantly different between groups at a false discovery rate of 0.05 were extracted for downstream analyses. A gene set was compiled containing genes associated with the Gene Ontology terms kinetochore, microtubule, mitotic spindle and mitosis.

### Statistical analyses

All data are shown as mean ± standard deviation (S.D.) or standard error of the mean (S.E.M.) as indicated. For comparisons, unpaired two-sided Student’s *t*-test or Fisher’s exact test were applied using the Prism software (GraphPad) as indicated. All cell-based *in vitro* experiments were independently repeated three times. No statistical method was used to pre-determine sample size throughout this study.

The investigators were not blinded to allocation for the *in vivo* experiments.

### Data availability

The HMECs and mouse tumor RNA sequencing data generated in this study will be deposited in a public database (will be available before publication). Secondary accessions: the human breast cancer RNA-sequencing data sets used to support the findings of this study are derived from the TCGA Research Network (http://cancergenome.nih.gov) and are available from GSE62944.

All other data supporting the findings of this study are available from the corresponding author upon request.

## Results

### MYC overexpression delays mitotic progression and causes CIN

We sought to determine if MYC alters mitotic progression in human non-transformed epithelial cell lines. We engineered human RPE-1 cells to constitutively overexpress MYC (RPE-MYC) or a control plasmid (RPE-NEO)^22,36^. Long-term MYC overexpression can lead to genomic changes and accumulation of mutations that might hinder interpretation. Thus, we also studied early passage non-immortalized human mammary epithelial cells (HMEC) that express a 4-hydroxytamoxifen (TAM)-activated *MYC-ER* transgene^37^; this allows for transient expression of MYC following treatment with TAM. We transduced cells with H2B-mCherry to follow chromosome alignment and segregation. Using time-lapse fluorescence microscopy, we determined the time from chromosome condensation in prophase to the onset of chromosome segregation in anaphase (**Figure 1**). MYC overexpression in RPE-1 cells markedly delayed anaphase onset. RPE-MYC cells required double the amount of time (58 minutes) from chromosome condensation to anaphase onset in comparison to 29 minutes for RPE-NEO (**Figure 1b, p <0.001**). Similarly, activation of MYC in HMECs (MYC ON) increased the time cells needed to proceed from prophase to anaphase from 24 to 36 minutes (**Figure 1b, p <0.001**). Increased time in prometaphase indicates prolonged SAC activation. To check if increased MYC activity causes SAC activation by inducing microtubule-kinetochore attachment errors we performed immunofluorescence of fixed cells (**Figure 1c**) to examine spindle morphology and chromosome localization. MYC activation was associated with increased appearance of metaphase cells with chromosomes located at the spindle pole (**Figure 1d**). Those chromosomes either take longer to align or they represent merotelic attachments. Merotelic chromosomes often cause lagging chromosomes in anaphase. Thus, we sought to determine if the frequency of lagging chromosomes is also increased in the presence of MYC overexpression. Indeed, increased MYC resulted in more cells with lagging chromosomes (**Figure 1d**). Often, lagging chromosomes are not incorporated into the nucleus, leading to the formation of micronuclei and CIN^38^. Consistently, increased micronuclei formation was observed upon expression of MYC in RPE-1 cells and activation of MYC in HMECs (**Figure 1d**). Thus, MYC overexpression or activation causes an increase in lagging chromosomes, micronuclei formation and CIN.

### MYC induced CIN is reversible

We next asked whether MYC-induced CIN is reversible in MYC-driven cancer cells. To test this, we derived a cell line from a MYC-driven transgenic mouse model of triple-negative breast cancer (MTB-TOM) in which MYC overexpression is inducible with doxycycline^37,39–41^. We isolated and immortalized MTB-TOM cells in presence of doxycycline to ensure MYC overexpression. Doxycycline-inducible MYC transgene expression was retained in MTB-TOM cells *in vitro* (**Figure 1e**), however, 2 days after withdrawing doxycycline, MYC expression is drastically reduced (**Figure 1e**). Importantly, in the absence of doxycycline, MTB-TOM derived tumor cells continue to proliferate, likely due to modest tonic level of endogenous murine MYC expression. We first determine if MTB-TOM cells grown in the presence of doxycycline (MYC ON) are chromosomally instable. Indeed, 15% of MYC overexpressing MTB-TOM cells have lagging chromosomes and 4% form micronuclei, a similar amount observed in RPE-MYC cells (**Figure 1f**). We next investigated whether turning off MYC expression would decrease CIN. Strikingly, the number of lagging chromosomes and micronuclei decreased to 1.2% and 1.3%, respectively (**Figure 1f**). We performed live cell fluorescence microscopy of H2B mCherry expressing MTB-TOM cells to determine whether the SAC is silenced earlier after MYC withdrawal. Indeed, MYC depletion caused a ~ 9 min decrease in the time cells require to go from prophase to anaphase, indicating an earlier satisfaction of the SAC (**Figure 1g**). We next wondered if attenuation of MYC expression could also reverse CIN in established human tumor cell lines. We quantified the number of micronuclei in two triple-negative breast cancer cell lines, HCC1143 and MDAMB231. Both cell lines were highly chromosomally unstable with 6.7% micronuclei detected in MDAMB231 and 21.5 % observed in HCC1143 cells (**Figure 1h**). Strikingly, reducing the amount of MYC by siRNA knock down reduced the number of cells with micronuclei significantly to 3.1 and 9.1 %. In summary, our data show that attenuation of MYC expression can reverse the appearance of lagging chromosomes, micronuclei and CIN. Thus, unlike MYC’s ability to accelerate transition through the G1/S checkpoint^42–44^, we find that increased MYC activity delays progression through mitosis and increases frequency of CIN in a reversible manner.

### MYC overexpression impairs mitotic spindle formation

Multiple kinds of mitotic errors can lead to merotelic attachments and CIN. We thus examined which steps of mitotic spindle assembly are affected by MYC overexpression. Brief treatment of cells with nocodozole followed by washout is a frequently utilized assay to analyze the timing of spindle re-assembly. We examined the timing of spindle formation including microtubule aster formation, centrosome movement and chromosome alignment in RPE-1 and HMECs. Strikingly, MYC overexpression interfered with numerous steps of spindle formation (**Figure 2a**). MYC most prominently affected microtubule nucleation and aster formation: MYC deregulation resulted in an increase in non-centrosomal microtubule nucleation (**Figure 2b and e**). Ten minutes after nocodazole washout microtubule asters formed at non-centrosomal sites in 100% of RPE-MYC cells compared to only ~ 50% of control cells (**Figure 2b**). Likewise, MYC activation in HMECs increased the percentage of cells with non-centrosomal microtubule asters from ~ 30 to ~ 80% after 10 minutes (**Figure 2e**).

**Figure 2.**
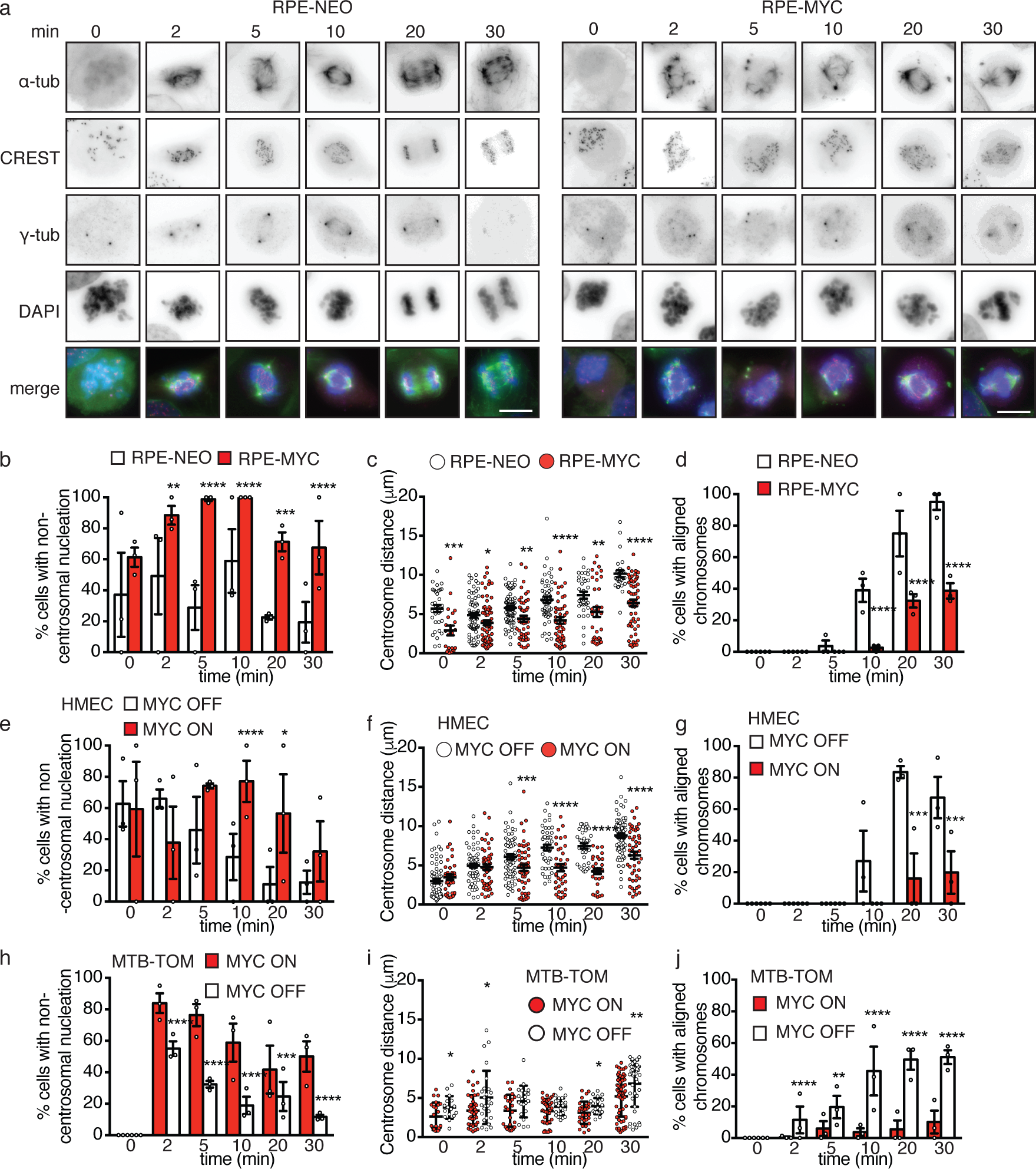
MYC influences microtubule nucleation, centrosome movement and spindle formation. **a** Representative images of a microtubule regrowth assay of RPE-NEO and RPE-MYC cells. Cells were fixed at indicated time points after nocodazole washout and stained with anti-δ-tubulin, anti-8-tubulin, anti-CREST and DAPI (scale bar, 10 μm). **b** Percent of RPE-NEO (white bars) and RPE-MYC (red bars) cells with non-centrosomal nucleation sites. **c** Centrosome distance in RPE-NEO (white circles) and RPE-MYC (red circles) **d** Percentage of RPE-NEO (white bars) and RPE-MYC (red bars) cells with aligned chromosomes. **e** Percent of HMECs expressing MYC-ER in the absence (MYC OFF, white bars) and presence of MYC activation for 3 days (MYC ON, red bars) with non-centrosomal nucleation sites. **f** Inter-centrosome distance in HMECs expressing MYC-ER in the absence (MYC OFF, white circles) and presence of MYC activation for 3 days (MYC ON, red circles) **g** Percentage of HMECs expressing MYC-ER in the absence (MYC OFF, white bars) and presence of MYC activation for 3 days (MYC ON, red bars) with aligned chromosomes. **h** Percent of MTB-TOM cells with non-centrosomal nucleation sites grown in presence (MYC ON, red bars) and absence (MYC OFF, white bars) of doxycycline for 4 days. **i** Centrosome distance of MTB-TOM cells grown in presence (MYC ON, red circles) and absence (MYC OFF, white circles) of doxycycline for 4 days. **j** Percentage of MTB-TOM cells with aligned chromosomes in presence (MYC ON, red bars) and absence (MYC OFF, white bars) of doxycycline for 4 days. Bars, mean +-S.E.M. n= 30-180 cells. **b, d, e, g, h, j** Fisher’s exact test. **c, f, i** t-test. * p<0.05, ** p< 0.01, *** p<0.001, **** p<0.0001.

Incomplete centrosome separation can lead to increased rates of chromosome missegregation^45,46^. We wondered whether MYC deregulation also affected centrosome movement. In control cells prior to nocodazole washout, centrosome positions were widely variable with a mean distance of 5.7 μm in RPE-NEO and 3 μmin HMECs MYC OFF (**Figure 2c and f**). After washout, centrosomes moved apart to reach a distance of 10.2 and 8.8 μm, respectively. However, only 38% of RPE-MYC cells established a centrosome distance of 9 μm or higher and in 38% of the cells centrosomes did not separate, indicated by a distance of less than 4 μm. (**Figure 2c**). Similarly, in HMEC cells treated with TAM (MYC ON) centrosome movement was slower and centrosome distance 30 minutes after nocodazole release was 2.4 μm shorter compared to HMEC-MYC cell not treated with TAM (MYC OFF) (**Figure 2f**). Finally, increased MYC led to a delay in chromosome alignment. 83% of RPE-NEO cells assembled a complete bipolar spindle with aligned chromosomes 30 minutes after washout, in contrast only 30% of RPE-MYC cells completed chromosome alignment in the same time frame (**Figure 2d**). A similar chromosome alignment delay was induced in HMECs after activation of MYC with 67% of control cells (MYC OFF) and only 20% of HMEC MYC ON cells aligned chromosomes 30 minutes after wash out (**Figure 2g**).

We next wondered if MYC depletion could rescue spindle formation defects and delays. We performed the nocodazole wash-out assay using the MTB-TOM cells derived from a primary MYC-driven breast tumor. Spindle assembly of MYC-overexpressing MTB-TOM cells (doxycycline treated) resembled the phenotype of RPE-MYC cells. Before removal of nocodazole centrosomes were positioned close together with a distance of 2.6 μm (**Figure 2i**). After washing out nocodazole many microtubules nucleated at the kinetochores (**Figure 2h**). Asters coalesced slowly into a bipolar spindle and only 10% of cells had aligned chromosomes 30 min after nocodazole washout (**Figure 2h and j**). Strikingly, switching off MYC expression rescued the spindle formation defects. Mean centrosome positioning before nocodazole removal increased from 2.6 to 3.8 μm (**Figure 2i**), the number of microtubule nucleation sites shortly after nocodazole wash out was reduced and nucleation was found predominantly at the centrosomes (**Figure 2h**). MYC depletion also improved chromosome alignment: 51% of cells aligned their chromosomes into a metaphase plate 30 minutes after nocodazole wash out in comparison to 10 % in the presence of high MYC (**Figure 2j**). Finally, reducing MYC expression increased the mean distance between centrosomes 30 minutes after washout from 5.2 to 6.8 μm (**Figure 2i**). Taken together, increased MYC influences several steps of mitotic progression including centrosome positioning, microtubule nucleation and growth resulting in a delay of chromosome alignment during mitosis. In contrast, attenuating MYC expression, even in established MYC-driven tumor cells, reverses these aberrant microtubule dynamics and improves chromosome alignment.

### MYC regulates genes involved in spindle formation

We next investigated how increased MYC activity might induce mitotic aberrations. MYC is a transcription factor that binds to perhaps thousands of promoters activating and repressing specific sets of direct target genes^47–50^. We used gene expression profiling to define the transcriptional effects of MYC on genes involved in mitotic progression. We analyzed the expression of 1452 genes associated with the gene ontology terms kinetochore, microtubule, mitosis and mitotic spindle (termed spindle genes). The expression of 416 spindle genes was significantly changed in HMEC treated with TAM (MYC ON) compared to control cells (MYC OFF) (**Supplementary table 1**). To check if spindle genes are also regulated by MYC in a tumor we analyzed gene expression of MTB-TOM tumors, in which MYC overexpression is induced by doxycycline (MYC ON) (**Figure 3a**). 456 spindle genes are differentially regulated in primary MTB-TOM tumors compared to normal mammary gland (**Supplementary table 1**). We extended our analysis to human breast cancer patient samples to see if MYC also influences the expression of spindle genes in a human cancer. Receptor triple-negative breast cancers (TNBCs) were previously found to express elevated MYC compared to receptor positive tumors^23,51^. We compared gene expression of triple-negative versus receptor positive breast cancers and excluded non-tumor tissues from our analysis to minimize the effects of proliferation on gene expression change. 179 spindle genes were differentially expressed in triple-negative, MYC high, compared to receptor positive, MYC low, tumors (**Supplementary table 1**). Taken together, the expression of at least 231 spindle genes were differentially regulated in 2 of 3 datasets and 34 genes were significantly deregulated in all 3 datasets (**Supplementary table 1, Figure 3b**). To further substantiate the influence of MYC we briefly turned off the expression of MYC in four MTB-TOM tumors by withdrawing doxycycline for 3 days (**Figure 3a**). MYC shutdown reversed the expression of approximately half of the MYC-induced spindle genes (124 of 231 genes) (**Figure 3c and Supplementary table 1 and 2**). We next examined the types of spindle genes that were deregulated by MYC by identifying those that could be grouped based on their known or suspected functions including: MT polymerization, MT depolymerization, motor proteins, centrosomal nucleation and acentrosomal nucleation (**Table 1**). We found that MYC altered the expression of genes in all four groups, however increased acentrosomal nucleation factors and motor proteins predominated. Interestingly, TPX2, BIRC5 and KIF11 regulate processes that we observed to be deregulated when MYC is overexpressed; TPX2 is the main factor that facilitates microtubule nucleation at chromosomes and its localization depends on BIRC5. TPX2 also regulates spindle pole segregation via KIF11. We found that the mRNA expression of each of these genes is induced by MYC and reversed upon MYC withdrawal in MTB-TOM tumors (**Figure 3d)**. In conclusion, MYC induces the expression of multiple genes involved in kinetochore function, microtubule biology and mitotic spindle formation; expression of many of these genes is reversible upon MYC withdrawal. We postulate that all or a subset of these MYC regulated genes could contribute to the observed MYC-dependent mitotic aberrations. We wondered whether the abnormalities observed in cells with increased MYC described above create new dependencies on the mitotic apparatus, which in turn might engender novel synthetic lethal interactions.

**Figure 3.**
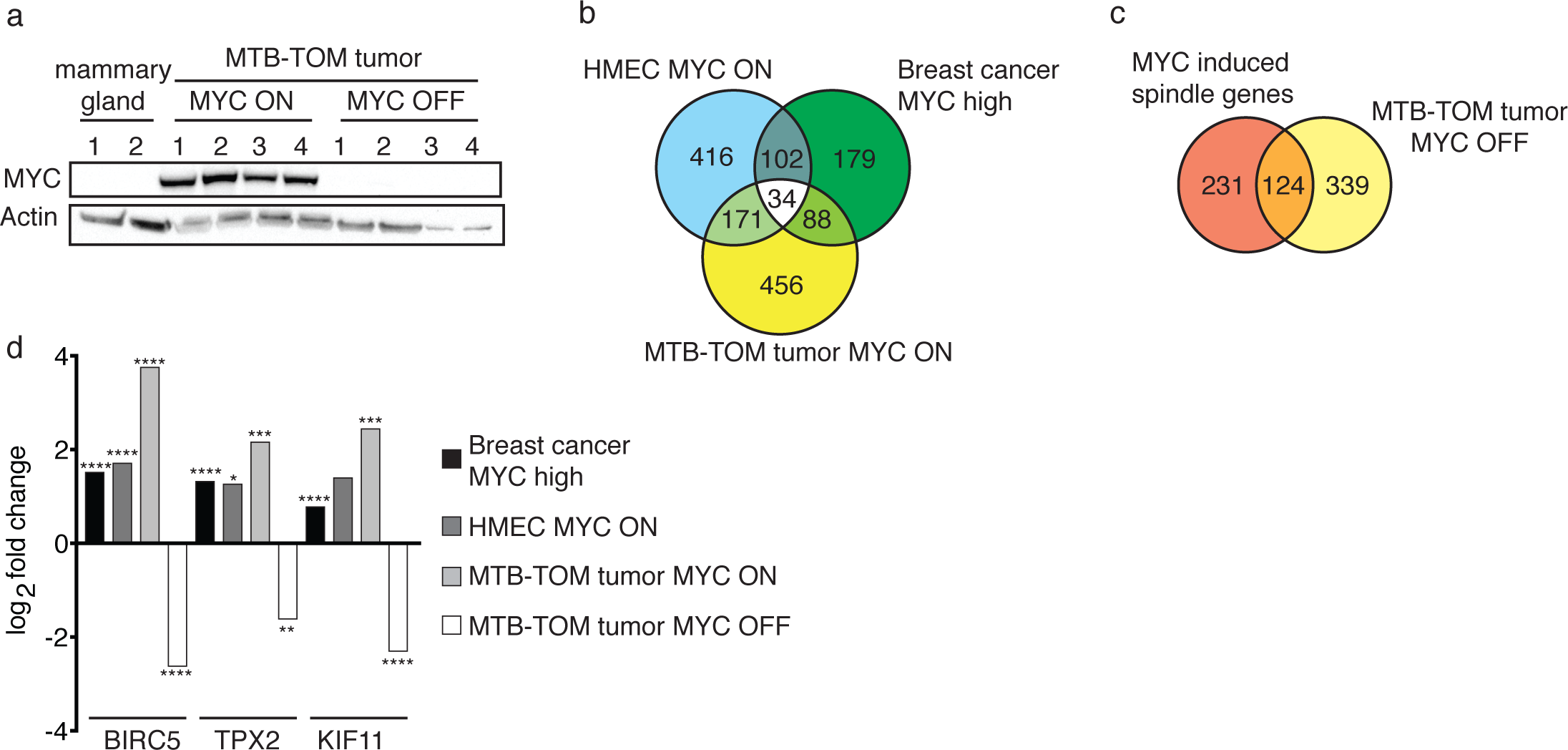
MYC regulates the expression of spindle genes. **a** Western blot of MYC in mouse mammary glands (n=2), MYC-driven tumors (MYC ON, n=4) and tumors from mice that were taken off doxycycline diet for 3 days to switch off MYC transgene expression (MYC OFF, n=4). **b** Overlap of spindle genes that are regulated by MYC in comparison to controls in the following three systems: HMECs in which MYC has been activated for 3 days (MYC ON) vs MYC-inactive HMECs, patient samples of MYC-high triple-negative breast cancer vs MYC-low receptor-positive breast cancer, and MYC-driven MTB-TOM tumors in the presence or absence of doxycycline (MYC-ON vs MYC-OFF, respectively). Genes differentially-regulated in at least 2 datasets were called MYC-induced spindle genes. **c** Overlap of MYC induced spindle genes with genes that are differentially regulated in MYC-driven mouse tumors in which MYC expression was switched off for 3 days. **d** Gene expression change of BIRC5, TPX2 and KIF11 in MYC high breast cancer (black bars), HMEC expressing MYC-ER in the presence of MYC activation (HMEC MYC ON, gray bars), MTB-TOM tumors (MYC ON, light gray bars) and MTB-TOM tumors in which MYC expression was switched off for 3 days (MYC OFF, white bars). * p<0.05, ** p< 0.01, *** p<0.001, **** p<0.0001.

**Table 1.** MYC regulates the expression of genes encoding for microtubule (MT) depolymerases, microtubule polymerases, motor proteins, proteins facilitating centrosomal nucleation and acentrosomal nucleation in HMEC-MYC, mouse MTB-TOM tumors (MYC ON), human triple-negative (MYC high) breast cancer tumors and mouse MTB-TOM tumors that were off doxycycline for 3 days (MYC OFF). Log2 fold change (logFC) and false discovery rate (FDR) are shown. Gray values: FDR>0.05.

|  | HMEC MYC ON |  | MTBTOM tumor MYC ON |  | Breast cancer MYC high |  | MTBTOM tumor MYC OFF |  |
| --- | --- | --- | --- | --- | --- | --- | --- | --- |
|  | Log2FC | FDR | Log2FC | FDR | Log2FC | FDR | Log2FC | FDR |
| **microtubule depolymerization** |  |  |  |  |  |  |  |  |
| KIF2C (MCAK) | 1.76 | >0.0001 | 2.78 | >0.0001 | 1.5 | >0.0001 | -1.86 | 0.0004 |
| STMN1 | 1.11 | 1.00000 | 2.41 | >0.0001 | 1.37 | >0.0001 | -1.73 | 0.0002 |
| **microtubule polymerization** |  |  |  |  |  |  |  |  |
| CKAP5 (ch-TOG) | 1.08 | 0.85 | -2.44 | 0.027 | 0.29 | >0.0001 | -0.47 | 0.32 |
| MAP4 | 1.13 | 0.32 | -1.28 | 0.04 | -0.05 | 0.31 | 1.22 | 0.02 |
| CENPE | -1.15 | 0.87 | 1.9 | 0.01 | 1.04 | >0.0001 | -1.55 | 0.02 |
| CLASP1 | -1.59 | >0.0001 | -0.40 | 0.44 | 0.21 | >0.0001 | 0.38 | 0.37 |
| **Motor proteins** |  |  |  |  |  |  |  |  |
| Kif15 | 1.47 | 0.04 | 1.96 | 0.0009 | 1.16 | >0.0001 | -2.07 | >0.0001 |
| KIF11 (Eg5) | 1.39 | 0.13 | 2.44 | 0.0007 | 0.78 | >0.0001 | -2.3 | 0.0002 |
| Kif22 | 1.42 | 0.0001 | 2.19 | >0.0001 | 0.06 | 0.42 | -1.73 | >0.0001 |
| KIFC1 | 2.2 | >0.0001 | 2 | 0.0001 | 1.34 | >0.0001 | -1.44 | 0.0015 |
| **centrosomal nucleation** |  |  |  |  |  |  |  |  |
| PLK1 | 1.69 | >0.0001 | 2.45 | 0.0002 | 1.39 | >0.0001 | -1.7 | 0.003 |
| AURKA | 1.38 | 0.01 | 4.18 | >0.0001 | 0.56 | >0.0001 | -2.69 | 0.0006 |
| TACC3 | 1.77 | 0.04 | 2.42 | >0.0001 | 0.92 | >0.0001 | -1.65 | 0.0002 |
| **acentrosomal nucleation** |  |  |  |  |  |  |  |  |
| TPX2 | 1.26 | 0.03 | 2.16 | 0.0008 | 1.32 | >0.0001 | -1.61 | 0.004 |
| DLGAP5 | 2.25 | >0.0001 | 2.19 | 0.0007 | 1.24 | >0.0001 | -1.52 | 0.007 |
| NUP107 | 1.46 | 0.001 | 1.42 | 0.001 | 0.18 | 0.002 | -1.48 | >0.0001 |
| NUSAP1 | 1.6 | >0.0001 | 1.84 | 0.004 | 0.71 | >0.0001 | -1.53 | 0.006 |
| TACC3 | 1.77 | 0.04 | 2.42 | >0.0001 | 0.92 | >0.0001 | -1.65 | 0.0002 |
| CDCA8 | 2.09 | >0.0001 | 2.87 | >0.0001 | 1.46 | >0.0001 | -2.18 | >0.0001 |
| INCENP | 1.65 | >0.0001 | 1.49 | 0.01 | 0.78 | >0.0001 | -1.29 | 0.01 |
| AURKB | 2.3 | >0.0001 | 2.93 | >0.0001 | 1.71 | >0.0001 | -2.46 | >0.0001 |
| BIRC5 | 1.7 | >0.0001 | 3.75 | >0.0001 | 1.51 | >0.0001 | -2.62 | >0.0001 |
| RANGAP1 | 1.45 | >0.0001 | 1.12 | 0.004 | 0.48 | >0.0001 | -1.21 | 0.0002 |
| LMNB2 | 1.33 | 0.0002 | 1.67 | >0.0001 | 0.88 | >0.0001 | -1.46 | >0.0001 |

### High TPX2 expression is required for mitotic completion in MYC-high cells

To test if cells with overexpressed MYC are dependent on TPX2, BIRC5 or KIF11, we performed siRNA knock down in RPE-NEO and RPE-MYC cells and measured cell death after 3 days (**Figure 4a**). We tested several control genes whose expression was not deregulated in a MYC dependent manner, but that are also important for mitotic function: MAP7, PDS5B and LMNA. We also examined Npmm1, whose expression is MYC-dependent, but is not directly involved in mitosis. As positive a control we included treatment with the cyclin-dependent kinase 1 (CDK1) inhibitor Purvalanol A, which has been previously shown to be synthetic-lethal with MYC overexpression^22^. Interestingly, the knock down of TPX2, BIRC5 and KIF11 markedly reduced the viability of RPE-MYC cells while minimally affecting the viability of RPE-NEO cells (**Figure 4a**).

Since TPX2 knock-down resulted in the greatest induction of cell death in MYC overexpressing cells, we sought to explore in more detail the connection between MYC and TPX2 expression. We found a significant co-occurrence of MYC and TPX2 mRNA upregulation in tumors of breast cancer patients (**cBioportal; Supplementary figure 1).** We next wondered if the MYC-TPX2 correlation we observed in the RNAseq datasets is also reflected in elevated TPX2 protein expression. Indeed, TPX2 protein correlates with increase MYC expression or activity in RPE-1 cells, HMECs, as well as in the MTB-TOM cell line (**Figure 4b,c and d**). We found rapid and reversible induction of TPX2 expression when MYC activity was regulated (**Fig 4c and d**), suggesting that MYC likely has a direct effect on TPX2 expression. We also observed that TPX2-MYC protein expression were correlated in breast cancer cell lines, with high levels of both proteins predominating in TNBC lines (**Figure 4e**). Likewise, in 12 patient-derived breast cancer xenograft models, TPX2 levels closely tracked MYC protein levels in TNBC models, but were less concordant in receptor positive (RP) PDX samples (**Figure 4f**). In the primary MYC-driven MTB-TOM tumor model, TPX2 protein level was increased compared to normal mammary gland tissue. Moreover, switching off MYC expression in the tumors reduced TPX2 RNA and protein to almost undetectable levels (**Figure 3d, 4g**). To determine if TPX2 is a transcriptional target of MYC we analyzed publically available Chip-seq data from the Encode project. MYC is found bound to the promoter region of TPX2 in multiple cell lines, suggesting that TPX2 is a direct transcriptional target of MYC (**Supplementary figure 2**). In a prior study, the TPX2 promoter was also found to be bound by MYC in murine fibroblasts, T-lymphoma cells and pancreatic tumor cells^50^, suggesting that MYC may drive TPX2 expression in multiple cell types and tumor contexts.

In non-transformed cells, TPX2 expression peaks in late G2 and mitosis and then TPX2 is degraded upon exit from mitosis, prior to the next cell cycle^52^. To examine where in the cell cycle TPX2 is expressed in control and MYC-high cells, we synchronized the RPE-NEO (control) or RPE-MYC cells. Cells were arrested at the G1/S boundary by a double thymidine block. After release (time 0) samples were taken every 2 hours for western blot analysis and cells observed by microscopy. Following release most RPE-NEO and RPE-MYC cells entered mitosis at ~ 10 hrs as evidenced by the observed cell rounding. However, more TPX2 protein is found in every cell cycle stage in RPE-MYC compared to RPE-NEO, demonstrating that the MYC-TPX2 correlation is not simply attributed to more cells in G2 or M phase in the setting of MYC overexpression (**Figure 4h**).

### TPX2 depletion is synthetic-lethal with MYC

In our screen, TPX2 depletion selectively caused cell death in RPE-MYC but not in RPE-NEO (**Figure 4a**). TPX2 knock down efficiency was subsequently confirmed by Western blot (**Figure 5a**). To further evaluate whether TPX2 induces apoptotic cell death only in RPE-MYC but not RPE-NEO cells, we monitored PARP cleavage. Notably, PARP cleavage is induced only in RPE-MYC and not RPE-NEO cells three days after TPX2 knock down (**Figure 5a**). Micrographs taken after treatment with control or TPX2 siRNA morphologic changes demonstrated cell death in RPE-MYC cells, but not RPE-NEO cells (**Figure 5b**). We next tested whether MYC-TPX2 synthetic-lethality is observed in other cell lines. Knock down of TPX2 in HMECs with MYC activation induced apoptotic PARP cleavage, but not in the control cells (**Figure 5c**). Bright field microscopy confirmed the selective cell death in HMECs with activated MYC following TPX2 knock down (**Figure 5d**). Next, we tested MYC-TPX2 synthetic lethality in breast cancer cell lines. We treated four triple-negative, high MYC and four receptor positive, low MYC breast cancer cell lines with control or TPX2 siRNA, and measured cell viability and PARP cleavage. Loss of TPX2 induced ~ 50 to 80% cell death in MYC-high triple-negative breast cancer cell lines (**Figure 5e**). However, the viability of only one receptor positive breast cancer cell line, T47D, decreased about 30% after TPX2 knock down, while the other RP cell lines were less sensitive. Similarly, PARP cleavage was observed in all triple-negative cell lines but only the T47D receptor positive cell line (**Figure 5f**). Despite being receptor positive, T47D cells have relatively high levels of MYC and TPX2, providing a possible explanation for the sensitivity of these cells to TPX2 depletion (**Figure 4c**).

**Figure 5.**
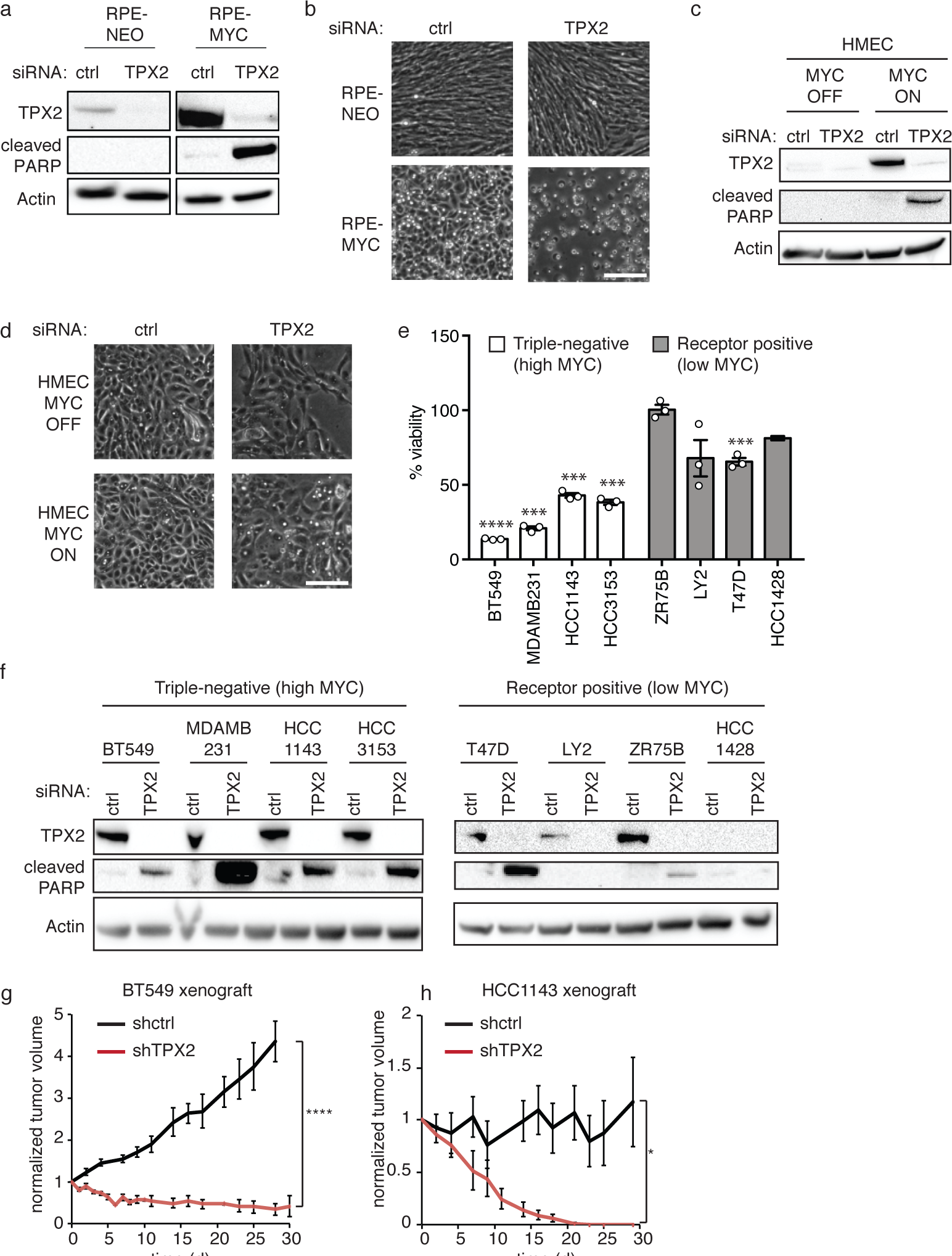
TPX2 is required for the survival of MYC high cells. **a** Western blot of TPX2 and cleaved PARP 3 days after transfection of RPE-NEO and RPE-MYC with control (ctrl) or TPX2 siRNA. **b** Micrographs of RPE-NEO and RPE-MYC 3 days after transfection with control (ctrl) or TPX2 siRNA. **c** Westernblot of TPX2 and cleaved PARP 3 days after transfection of HMEC expressing MYC-ER in the absence (MYC OFF) or presence of MYC activation (MYC ON) for 3 days with control (ctrl) or TPX2 siRNA. **d** Micrographs 3 days after transfection with control (ctrl) or TPX2 siRNA of HMEC expressing MYC-ER in the absence (MYC OFF) or presence of MYC activation for 3 days (MYC ON). **e** Percent viability of triple-negative, high MYC, and receptor positive, low MYC, breast cancer cell lines 3 days after transfection with TPX2 siRNA normalized to control siRNA. Bars, mean +-S.E.M., t-test, n=3. *** p<0.001, **** p<0.0001. **f** Western blot of TPX2 and cleaved PARP in triple-negative and receptor positive breast cancer cell lines 3 days after transfection with control (ctrl) or TPX2 siRNA. **g** and **h** Relative xenograft tumor volume of the triple-negative breast cancer cell lines BT549 (**g**) and HCC1143 (**h**). Cells expressed doxycycline inducible shRNA against TPX2 (shTPX2) or GFP (shctrl); doxycycline administration induced shRNA expression. Tumor volume was normalized to the volume at the beginning of shRNA induction. Mean +-S.E.M. BT549 shctrl (n=7), shTPX2 (n=7) HCC1143 shctrl (n=6), HCC1143 shTPX2 (n=5). t-test. * p<0.05, **** p<0.0001.

The most stringent test of a synthetic-lethal interaction is to determine if it can block *in vivo* tumor formation. Since a selective small molecule inhibitor of TPX2 does not presently exist, we sought to test if TPX2 depletion can regress xenograft tumors from MYC-high TNBCs. We generated two triple-negative breast cancer cell lines (BT549 and HCC1143) that express a doxycycline inducible shRNA against TPX2 (shTPX2) or GFP as control (shctrl). Importantly, incubation with doxycycline induces TPX2 knock down and cell death *in vitro* (**Supplemental Figure 2a and b**). We orthotopically transplanted shTPX2 or shctrl cells into the mammary fad pad of immunocompromised mice. Once the tumor reached 1 cm in greatest diameter, doxycycline was administered to induce shTPX2 or shctrl expression. BT549 tumors shrank markedly upon expression of shTPX2 (**Figure 5g**). Consistently, HCC1143 tumors completely disappeared 20 days after the induction of shTPX2 expression, indicating that TPX2 is required for tumor survival *in vivo* (**Figure 5h**). Tumor growth was not affected by shctrl expression. Taken together, these data demonstrate that TPX2 is a novel synthetic-lethal interaction partner of MYC.

### TPX2 expression is required for spindle assembly in MYC-high cells

TPX2 is found in the nucleus in interphase and localizes to spindle pole microtubules upon nuclear envelope breakdown in early mitosis^52^. We sought to determine if TPX2 localization is altered in RPE-MYC compared to RPE-NEO cells using immunofluorescence. TPX2 localization did not change upon expression of MYC in RPE-1 cells (**Supplementary Figure 3**). TPX2 knock down causes spindle formation failure and subsequent cell cycle arrest in Hela cells, that reportedly express high levels of MYC^52–54^. However, the consequences of TPX2 knock down in non-transformed, MYC-low expressing cells has not been reported. To investigate whether RPE-MYC cells die in the absence of TPX2 because of mitotic failure, we performed live cell imaging of RPE-NEO and RPE-MYC cells expressing the FUCCI cell cycle reporter^55^ (**Figure 6a**). After TPX2 knock down the majority (55%) of RPE-MYC cells died in mitosis after a prolonged arrest. An additional 23% of the arrested cells slipped into G1 and subsequently died, indicative of mitotic catastrophe. 48 hours after siRNA treatment only 20% of RPE-MYC cells were viable (**Figure 6b**). In contrast, only 8% of RPE-NEO cells died during the time course of the experiment (48 hrs). However, fewer RPE-NEO cells entered mitosis (64 cells entered mitosis after TPX2 KD in comparison to 159 cells treated with control siRNA), indicating partial cell cycle arrest at the G2**/**M checkpoint (**Figure 6b; last column**). We performed DNA content analysis 3 days after TPX2 knock down and confirmed a partial G2/M arrest in RPE-NEO cells while RPE-MYC cells underwent dramatic cell death (**Figure 6c**). We observed 43% apoptotic cell death in RPE-MYC cells, indicated by a smaller than 2N DNA content, compared to 15% in RPE-NEO cells. Furthermore, 25% of RPE-MYC cells contained a larger than 4N DNA content supporting the previous observed mitotic slippage. To understand if RPE-MYC cells could not proceed through mitosis after TPX2 knock down because of mitotic spindle formation failure we fixed and stained for components of the mitotic spindle (**Figure 6d**). TPX2 knock down drastically affected spindle formation in RPE-MYC cells. We found that 84% of RPE-MYC cells did not form a spindle in comparison to only 43% of RPE-NEO (**Figure 6e**). In contrast, 45% of RPE-NEO cells were able to form either normal-appearing or small spindles. TPX2 depletion also affected centrosome integrity differently in RPE-NEO and RPE-MYC. After TPX2 knock down, extra γ-tubulin positive microtubule foci appeared in both RPE-NEO and RPE-MYC as previously noted^56,57^ (**Figure 6d arrows**). However, in 60% of RPE-MYC cells none or only 1 γ-tubulin signal was detected. Loss of γ-tubulin foci was accompanied by a loss of microtubule asters, as 24% of RPE-MYC cells had less than 2 microtubule asters (**Figure 6f**). In contrast, TPX2 knock down in RPE-NEO did not reduce the number of γ-tubulin foci and microtubule asters. Our data suggest that TPX2’s function in recruiting γ-tubulin to the centrosome is critical for MYC high cells to nucleate microtubules and to form the mitotic spindle. These results indicate that non-transformed cells are less sensitive to loss of TPX2 and very low levels of TPX2 are sufficient for spindle assembly and progression through mitosis.

**Figure 6.**
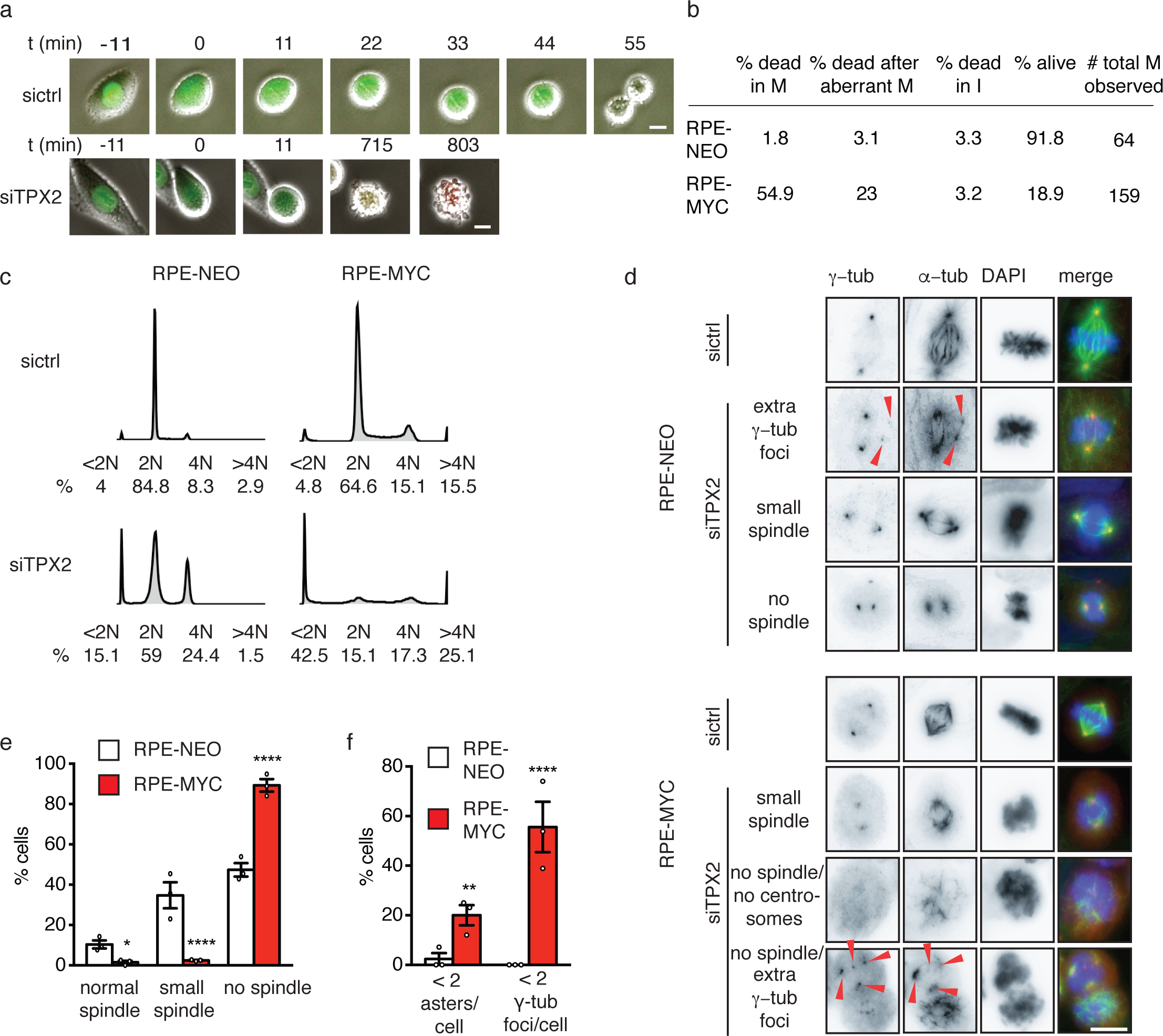
TPX2 protects the mitotic spindle function in MYC high cells. **a** Representative time-lapse microscopy images of RPE-MYC cells expressing the FUCCI cell cycle marker 12 hours after transfection with control (sictrl) or TPX2 (siTPX2) siRNA. Fluorescence and phase contrast images were taken every 11 min and overlaid. Scale bar, 20 μm. **b** Quantification of cells undergoing cell death in mitosis (M), after aberrant mitosis, in interphase (I) and number (#) of total mitotic cells observed 12 to 24 hours after TPX2 knock down in RPE-NEO and RPE-MYC. n=3. **c** Cell cycle profiles of RPE-NEO and RPE-MYC cells 48 hours after treatment with control (sictrl) or TPX2 (siTPX2) siRNA **d** Representative images of RPE-NEO (top) and RPE-MYC (bottom) 18 hours after transfection with control (sictrl) or TPX2 (siTPX2) siRNA. Cells were fixed and stained with α-tubulin, γ-tubulin and DAPI. After TPX2 knock down cells formed extra γ-tubulin foci, small spindles or no spindles. Some RPE-MYC cells did not exhibit any γ-tubulin foci. **e** Quantification of RPE-NEO and RPE-MYC cells with normal, small or no spindles 24 hours after transfection with TPX2 siRNA (siTPX2). **f** Percent of RPE-NEO and RPE-MYC with less than 2 microtubule asters and γ-tubulin foci per cell 24 hours after TPX2 knock down. **e and f** Bars, mean +-S.E.M, Fisher’s exact test, n=36 and 126. * p<0.05, ** p<0.01, **** p<0.0001.

### High TPX2 levels are protective for cells that overexpress MYC

Exogenous TPX2 overexpression leads to defects in microtubule organization and the formation of aberrant mitotic spindles^52,58^. We wondered whether the highly increased TPX2 expression found in RPE-MYC cells causes spindle formation defects, or rather if increased TPX2 is requisite for MYC-high cell survival. We tested several doxycycline-inducible shRNAs against TPX2 in RPE-MYC cells and identified one that partially lowers the amount of TPX2 to a level found in RPE-NEO cells (**Figure 7a)**. Partial depletion of TPX2 in RPE-MYC cells did not induce cell death **(Figure 7b)**, rather cells were further delayed in mitosis (mean time in mitosis increased from 40 min to 67 min) upon induction of shTPX2 (**Figure 7c**). Partial TPX2 depletion did not rescue the increased number of micronuclei found in RPE-MYC cells but further increased the number of micronuclei from 7.2 % found in control cells to 13.7 *%* in cells with shTPX2 expression (**Figure 7d**). We next performed the spindle re-assembly after nocodazole wash out assay to determine if any of the other MYC induced aberrations are rescued (**Figure 7e**). Lowering TPX2 did not decrease the number of microtubule nucleation sites at chromosomes (**Figure 7f**). Nor did it enhance the time cells need to align their chromosomes into a metaphase plate (**Figure 7g**). In fact, it further slowed chromosome alignment as 90 min after nocodazole wash out less than 5% of cells demonstrated aligned chromosomes (**Figure 7e,g**). Lowering TPX2 also slowed centrosome movement; even 90 mins after nocodazole wash out cells with partial depletion of TPX2 had shorter media inter-centrosome distance (**Figure 7h**).

**Figure 7.**
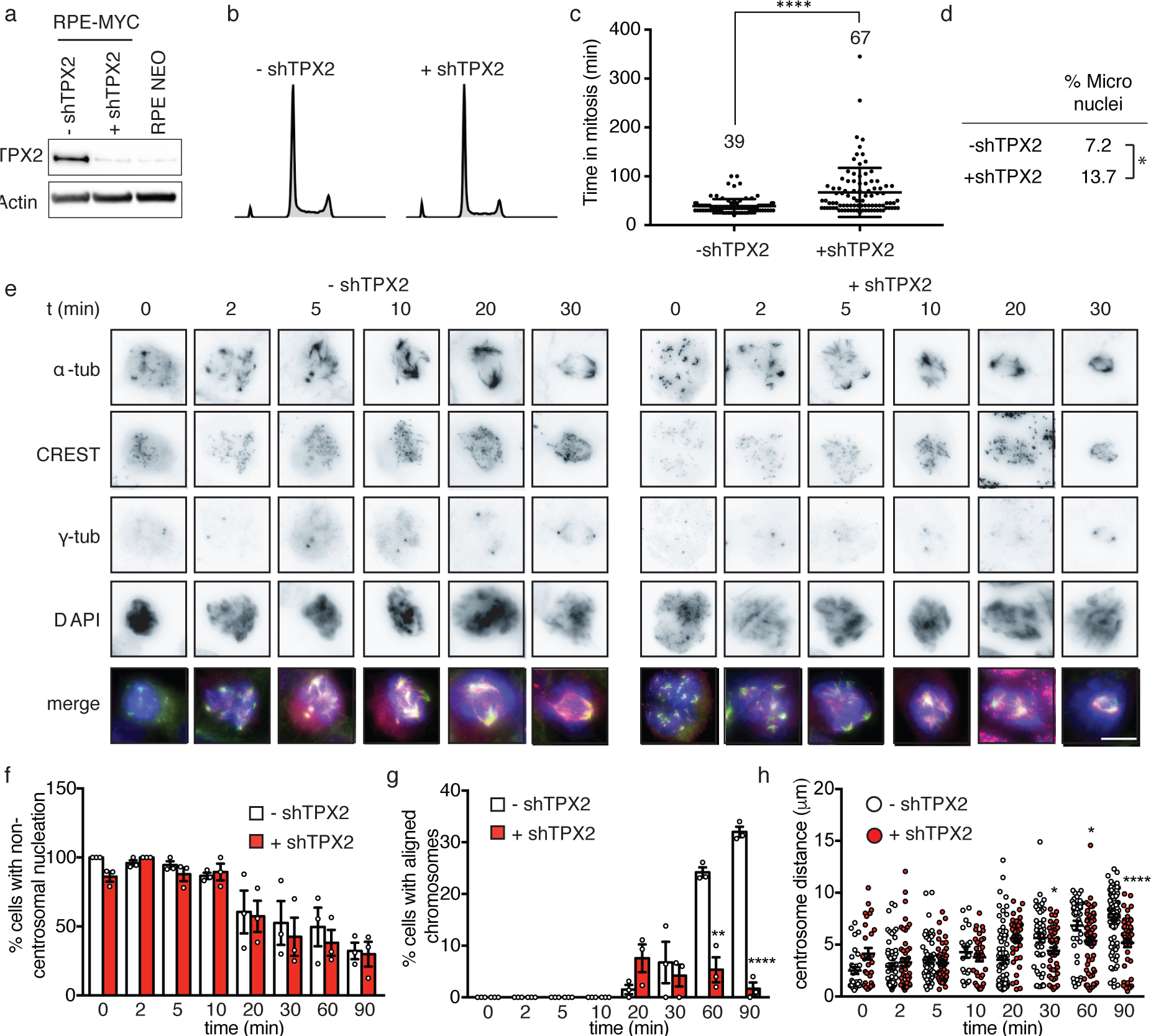
High levels of TPX2 are necessary for MYC high cells to progress through mitosis. **a** Westernblot of TPX2 in RPE-NEO and RPE-MYC expressing doxycycline inducible shRNA against TPX2 in absence (-shTPX2) and presence (+shTPX2) of doxycycline for 4 days. **b** Cell cycle profiles of RPE-MYC expressing doxycycline inducible shRNA against TPX2 in absence (-shTPX2) and presence (+shTPX2) of doxycycline for 4 days. **c** Quantification of time from nuclear envelop breakdown to anaphase onset from time-lapse microscopy of RPE-MYC expressing doxycycline inducible shRNA against TPX2 in absence (-shTPX2) and presence (+shTPX2) of doxycycline for 4 days. Mean +-S.D., t-test, n=164. **** p<0.0001. **d** Quantification of micronuclei of expressing doxycycline inducible shRNA against TPX2 in absence (-shTPX2) and presence (+shTPX2) of doxycycline for 4 days. Fisher’s exact test, n=223 and 182. * p<0.05. **e** Representative images of microtubule regrowth assay of expressing doxycycline inducible shRNA against TPX2 in absence (-shTPX2) and presence (+shTPX2) of doxycycline for 4 days. Cells were fixed at indicated time points after nocodazole washout and stained with anti-α-tubulin, anti-δ-tubulin, anti-centrosome and DAPI, scale bar, 10 μm. **f** Percent non-centrosomal nucleation sites in RPE-MYC expressing doxycycline inducible shRNA against TPX2 in absence (-shTPX2, white bars) and presence (+shTPX2, red bars) of doxycycline for 4 days. Bars, mean +-S.E.M, Fisher’s exact test, n=150-300. * p<0.05, ** p< 0.01, **** p<0.0001. **g** Percentage of aligned chromosomes in RPE-MYC expressing doxycycline inducible shRNA against TPX2 in absence (-shTPX2, white bars) and presence (+shTPX2, red bars) of doxycycline for 4 days. **h** Centrosome distance over time in RPE-MYC expressing doxycycline inducible shRNA against TPX2 in absence (-shTPX2, white bars) and presence (+shTPX2, red bars) of doxycycline for 4 days. Bars, mean +-S.E.M. **f and g** Fisher’s exact test, n=150-300. * p<0.05, ** p< 0.01, **** p<0.0001. **h** t-test n= 20-84 cells.

In summary, lowering TPX2 to levels found in non-transformed cells does not rescue the effects of MYC overexpression on mitotic progression and chromosome segregation, but rather further accentuates these defects. We thus conclude that high TPX2 expression partially rescues MYC induced mitotic defects and permits cells to proceed through mitosis.

## Discussion

MYC overexpression in tumors has been associated with aneuploidy, however whether MYC directly induces aneuploidy by eliciting CIN was not known. Here we show that MYC overexpression directly and reversibly induces chromosome missegregation and CIN by impacting multiple mitotic steps including: microtubule nucleation, microtubule aster coalescence and centrosome movement. We find that MYC overexpression changes the expression of multiple spindle-related genes, all of which might contribute to the observed spindle abnormalities. The net effects of these alterations, we show herein, is to alter spindle and spindle-pole morphology and delay progression through mitosis.

We found in multiple cell lines, in mouse tumor models and in patient samples of human cancer that cells with deregulate MYC expression rely on TPX2 overexpression to successfully complete mitosis. TPX2 plays a protective role by limiting the accumulation of mitotic abnormalities and sufficient CIN to elicit mitotic arrest and subsequent cell death. Depletion of TPX2 reveals a role in centrosome maturation and microtubule nucleation that is particularly important for MYC high cells. Centrosomes, while not essential, are the major sites of microtubule nucleation during spindle formation. Our work raises the possibility that overexpression of MYC impairs centrosome function, thus cells become reliant on acentrosomal mechanisms of spindle formation. TPX2’s major role in acentrosomal microtubule nucleation might explain the addiction of cells with deregulated MYC to high levels of TPX2.

Several mitotic functions of TPX2 are mediated by its role as an Aurora kinase A (AURKA) activator^59–61^. AURKA inhibition is synthetic-lethal with MYC, however, MYC degradation in interphase rather than a mitotic role has been proposed to cause cell death after AURKA inhibition^24,62^. Other functions of TPX2, besides AURKA activation, are likely important for the observed MYC-TPX2 dependency. In our screen, we identified two TPX2 interacting proteins, BIRC5 and KIF11, which are also required for the survival of cells with deregulated MYC. TPX2 recruits KIF11 to microtubules where it regulates microtubule density and spindle length. TPX2 can act as a scaffold protein for the chromosomal passenger complex where it interacts with BIRC5 and activates AURKB^63–65^. Thus, further studies are required to elucidate which of the multiple functions of TPX2 are required to attenuate MYC-induced mitotic spindle errors.

TPX2 is overexpressed in many cancers and was one of the genes whose expression most strongly correlates with a CIN phenotype across multiple different tumor types^34,35,66^. In addition to MYC, other tumor suppressor genes or oncogenes have been shown to induce CIN through mitotic defects^67,68^. Perhaps high levels of TPX2 might be protective for various types of cancers to prevent the development of physiologically intolerable levels of CIN, providing a possible explanation for the abundant upregulation of TPX2 in multiple cancers.

Finally, the reversibility of the observed MYC-induced spindle abnormalities and CIN indicates that MYC overexpression not only initiates CIN but is also required for its maintenance. Moreover, increased MYC expression has been observed in early tumor metastasis^69^ (Lawson, 2015) and drug resistant cancers^70^, situations associated with CIN and tumor evolution. Several small molecule inhibitors that can modulate MYC transcriptional activity are currently under preclinical development and evaluation^71–73^. We postulate that inhibition of MYC and thereby CIN might be a useful strategy to prevent tumor evolution, drug resistance and subsequent tumor relapse. Since TPX2 protein structure is largely intrinsically disordered, the development of anti-cancer drugs to directly inhibit TPX2 activity may pose a challenge^74^. However, inhibitors against the KIF11 motor and other mitotic proteins, potential synthetic-lethal interactors with MYC, are available and are currently being tested in clinical trials^75,76^. We propose that the expression of MYC and TPX2 in tumors might be useful biomarkers to stratify patients for new anti-mitotic therapies.

## Author Contributions

All authors listed, have made substantial, direct and intellectual contribution to the work, and approved it for publication. J.R. and A.G. conceived of the project, designed experiments and wrote the manuscript. J.R. led the research efforts and carried out most of the studies. A.C. performed siRNA knock-down studies; MT helped with microtubule regrowth assays; A.N.C., S.K.A., R.C. and S.B. assisted with RNAseq and bioinformatics. M.J. helped with TPX2 cell cycle distribution. A.G. supervised studies.

## Acknowledgements

We thank Dr. Wittmann and Dr. van Haren for plasmids and help with microscopy. We thank Dr. JM Bishop for cell lines, and Dr. Stephen Floor for help with Encode Analysis. We thank Dr. Sophie Dumont and all the members of the Goga laboratory for critically reading the manuscript. This work was supported by the Susan G Komen postdoctoral fellowship grant PDF15331114 (J.R.), a CIRM Predoctoral Award (M.T), the US NIH F99/K00 Predoctoral to Postdoctoral Transition Award F99CA212488 (R.C.), the CDMRP Breast Cancer Research Program (W81XWH-12-1-0272) (A.G.), NIH R01-CA170447 (A.G.) and Lymphoma Scholar Award (A.G.).

---

**Supplementary figure 1.**
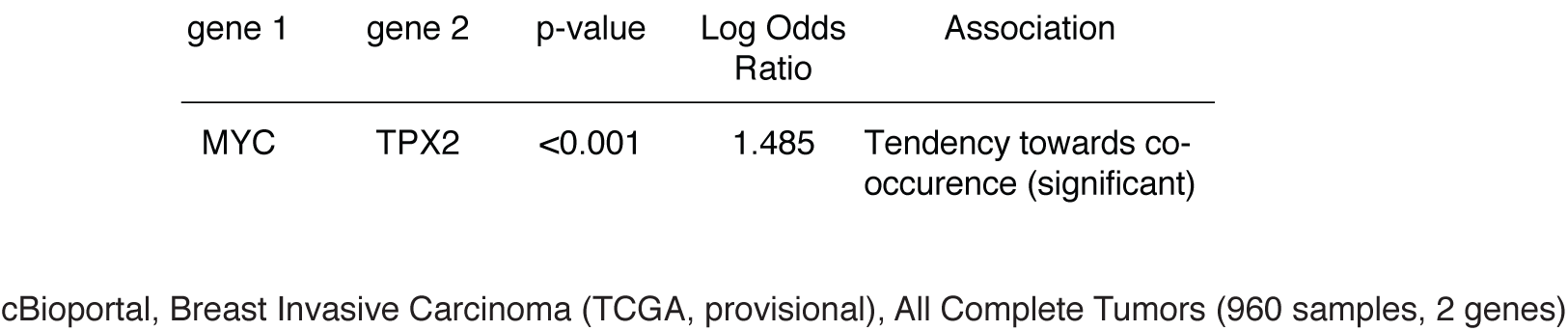
MYC and TPX2 mRNA expression correlate in human breast cancer. Data were obtained from cBioportal, Breast Invasive Carcinoma (TCGA, provisional).

**Supplementary figure 2.**
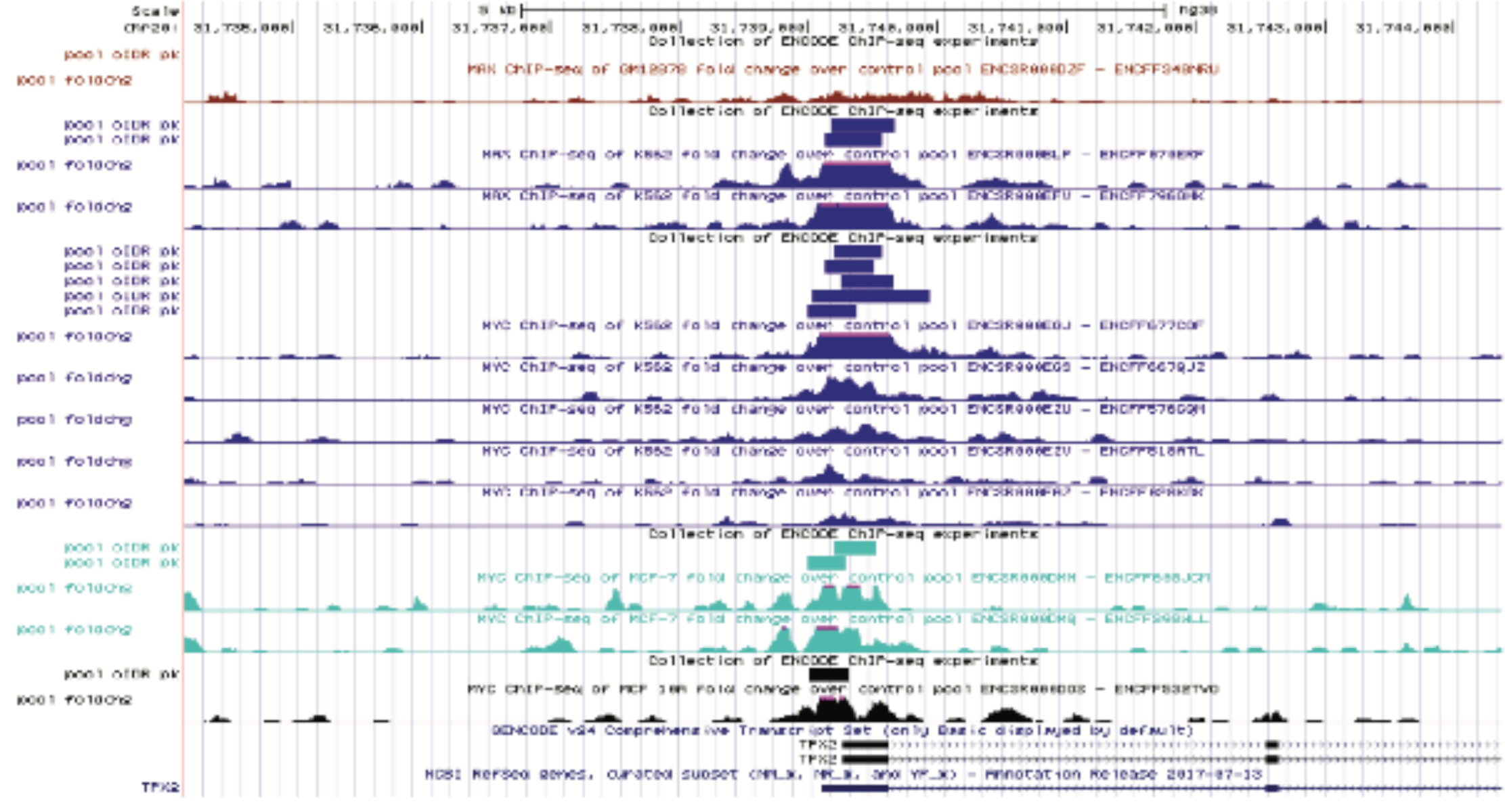
MYC/MAX bind to the promoter region of TPX2. ChIP seq data for MYC and MAX binding to the promoter region of TPX2 (chr20:31,734,166-31,744,376) from the ENCODE project (human Dec 2013 (GRCh38/hg38) assembly).

**Supplementary figure 3.**
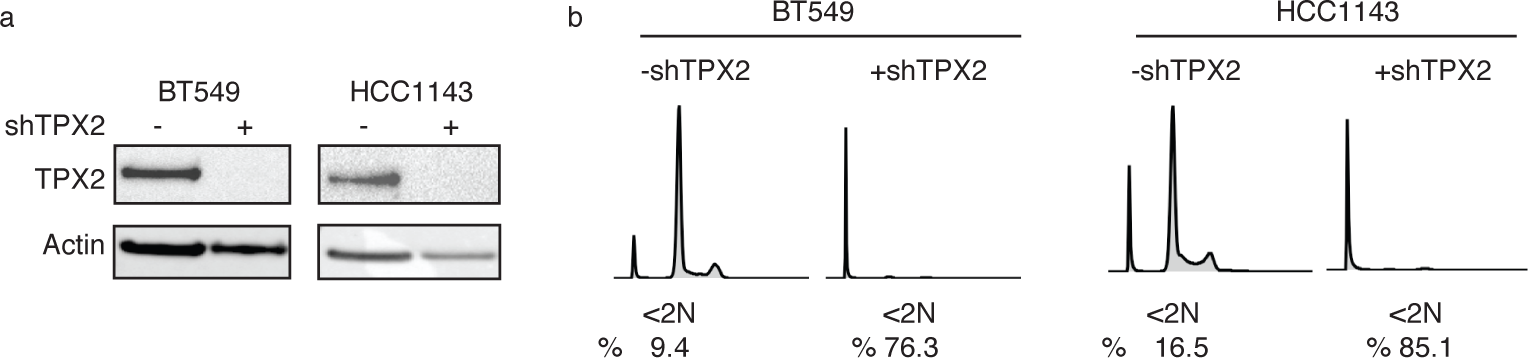
**a** Western blot for TPX2 in the triple-negative breast cancer cell lines BT549 and HCC1143 expressing a doxycycline-inducible shRNA against TPX2 in absence and presence of doxycycline for 5 days. **b** Cell cycle profile of BT549 and HCC1143 expressing a doxycycline-inducible shRNA against TPX2 in absence and presence of doxycycline for 5 days. Percent sub-2N DNA content is indicated.

**Supplementary figure 4.**
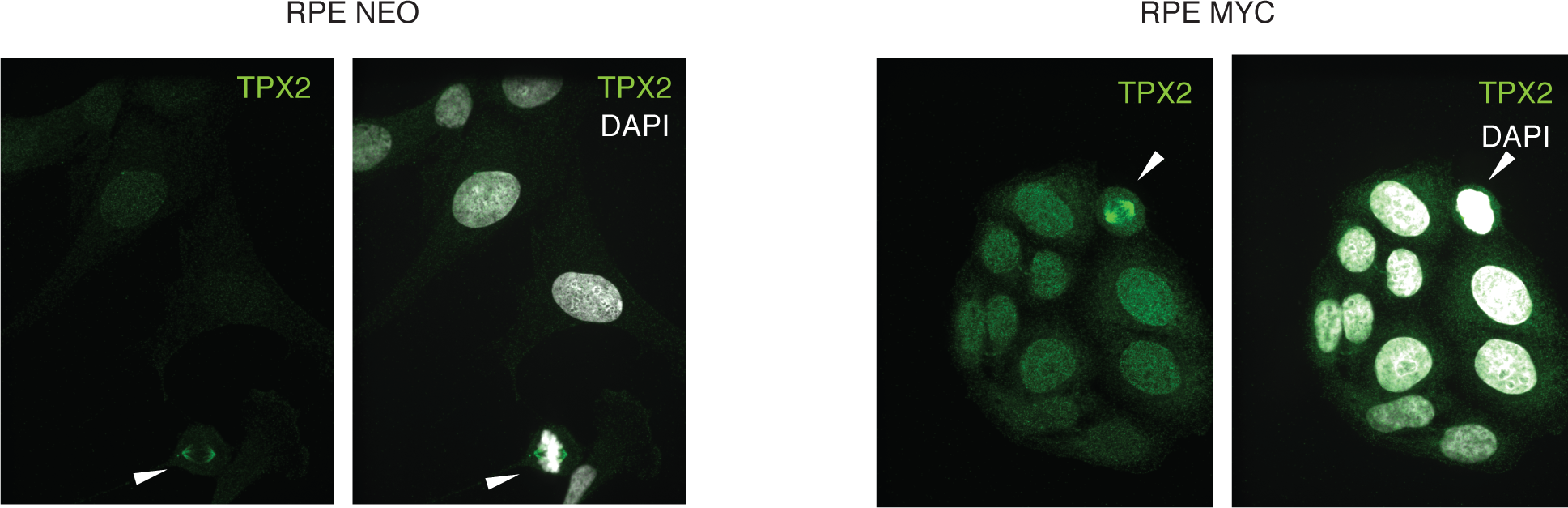
RPE-NEO and RPE-MYC cells were fixed and stained with an antibody against TPX2 (green) and DAPI (white).

**Supplementary table 1** Gene expression data from HMEC-MYC, mouse MTB-TOM tumors MYC ON and human triple-negative (MYC high) breast cancer tumors and mouse MTB-TOM tumors that were off doxycycline (MYC OFF) for 3 days to switch off MYC expression. Genes with a log2 fold change (logFC) > 1 and false discovery rate (FDR) <0.05 are shown.

**Supplementary table 2** Genes whose expression changed in HMEC-MYC, mouse MTB-TOM tumors MYC ON and human triple-negative (MYC high) breast cancer tumors, 2 of the 3 datasets (spindle genes) and spindle genes whose expression were reversed in MTB-TOM MYC OFF are listed. Log2 fold change (logFC) > 1, false discovery rate (FDR) <0.05.

